# The Pleiades cluster of fungal effector genes inhibit host defenses

**DOI:** 10.1101/827600

**Authors:** Fernando Navarrete, Nenad Grujic, Alexandra Stirnberg, David Aleksza, Michelle Gallei, Hazem Adi, Janos Bindics, Marco Trujillo, Armin Djamei

**Affiliations:** Gregor Mendel Institute (GMI), Austrian Academy of Sciences (OEAW), Vienna BioCenter (VBC), Dr. Bohr-Gasse 3, 1030 Vienna. Austria; Universität für Bodenkultur Wien. Department for Agrobiotechnology (IFA-Tulln). Konrad Lorenz Strasse 20. 3430 Tulln a.d. Donau. Austria; Institute of Science and Technology Austria, Am Campus 1, 3400 Klosterneuburg.; Institute of Molecular Biotechnology, Dr. Bohr-Gasse 3, 1030 Vienna. Austria; Albert-Ludwigs-University Freiburg, Faculty of Biology, Institute of Biology II, 79104 Freiburg, Germany; Leibniz Institute of Plant Genetics and Crop Plant Research (IPK), OT Gatersleben, Corrensstraße 3, D-06466 Stadt Seeland, Germany

**Keywords:** gene cluster, paralogs, effectors, *Ustilago maydis*, filamentous fungus, plant pathogen, virulence factor, maize, PAMP-triggered immunity, Reactive Oxygen Species, E3 ligase

## Abstract

**Summary:** Biotrophic plant pathogens secrete effector proteins to manipulate the host physiology. Effectors suppress defenses and induce an environment favorable to disease development. Sequence-based prediction of effector function is difficulted by their rapid evolution rate. In the maize pathogen Ustilago maydis, effector-coding genes frequently organize in clusters. Here we describe the functional characterization of the pleiades, a cluster of ten symplastic effectors. Deletion of the pleiades leads to strongly impaired virulence and accumulation of reactive oxygen species (ROS) in infected tissue. Eight of the Pleiades suppress the production of ROS upon perception of pathogen associated molecular patterns (PAMPs). Although genetically redundant, the Pleiades target different host components. The paralogs Taygeta1 and Merope1 suppress ROS production in either the cytoplasm or nucleus, respectively. Merope1 targets and promotes the autoubiquitination activity of RFI2, a conserved family of E3 ligases that regulates the production of PAMP-triggered ROS burst and influences flowering time in plants.

## Introduction

Plants are constantly colonized by a myriad of microbial organisms, yet for the most part, they remain asymptomatic. In order to interact with their host plants, pathogenic microbes evolved molecules known as effectors. These secreted molecules (proteins, RNA and small metabolites) manipulate host physiology and development to suppress immune responses and create an environment that promotes the pathogen’s proliferation. Following secretion, effector molecules can either remain in the space between the plant cells, the apoplast (apoplastic effectors) or be translocated into the host cytosol (symplastic effectors) (Sánchez-Vallet et al., 2018; Uhse & Djamei, 2018). As a result of being secreted molecules, effectors are exposed to the host immune system and can induce resistance responses following their recognition. Hence, the evolution of effector coding genes is governed by two major processes, evasion of host recognition and functional optimization, both of which can result in gene deletion events, gain of new effectors, point mutations, or alteration of gene expression (Sánchez-Vallet et al., 2018).

A major problem in effector biology is how to assign function to genes encoding putative effectors. Typically, the genomes of filamentous pathogens code for hundreds or even thousands of effector candidate genes (Duplessis et al., 2011; Lo Presti et al., 2015; Sánchez-Vallet et al., 2018) yet, for any given pathogen, only a handful have been functionally characterized. Predicting effector gene function from their sequence is hampered by the fact that most of them lack conserved protein domains or homologs beyond closely related species (Schirawski et al., 2010). They are also difficult to study by reverse genetic approaches as they generally exhibit functional redundancy. Large scale single knock out studies have identified only a few genes with a measurable contribution to virulence (Hiromasa Saitoh 2012; Uhse et al., 2018). This redundancy is believed to counteract recognition by the host immune system. If a single effector targets a given host pathway, losing this effector to avoid host recognition will cause a fitness cost to the pathogen population. However, if multiple effectors target the same pathway, the population can quickly adapt simply by losing the recognized effector without a fitness cost (Sánchez-Vallet et al., 2018). For example, many fungal pathogens have multiple LysM effectors that bind chitin in order to avoid host recognition or shield their cell wall from degradation by chitinases (de Jonge et al., 2010; Koharudin et al., 2011; Marshall et al., 2011). Another example is *Phytophthora infestans*, which secretes two protease inhibitors targeting the same host protease (Song et al., 2009).

In many genomes of plant interacting microbes, effector encoding genes are enriched in discrete regions of the genome and these regions show a higher sequence diversity within the population in comparison to housekeeping genes (Dong, Raffaele, & Kamoun, 2015; Sánchez-Vallet et al., 2018). An extreme case of genome compartmentalization is that of accessory or supernumerary chromosomes as in *Fusarium oxysporum* f. sp. *lycopersici* and *Zymoseptoria tritici*. These are polymorphic chromosomes not shared by all members of a population, rich in effector genes and dispensable for growth in axenic culture. Additionally, these genomic regions are enriched in transposon sequences, have low gene density, are located to sub-telomeric parts of the chromosomes and/or frequently show presence/absence polymorphisms. All of these features are believed to increase the variability of the effector repertoire in a pathogen population (Sánchez-Vallet et al., 2018).

Smut fungi are a class of fungal pathogens that infect grasses and some dicotyledonous plants, producing high amounts of spores in the floral tissues (Kamper et al., 2006; Sharma, Mishra, Runge, & Thines, 2014). *Ustilago maydis* infects maize and teosintes, inducing the production of galls in all aerial tissues as soon as 7 days post infection. This feature, together with its genetic tractability, has established the *U. maydis*-maize pathosystem as a model to study biotrophic plant pathogen interactions. In *U. maydis*, as in other smuts, a large proportion of effector genes are organized in clusters. These effector clusters consist of groups of 3 to 26 genes in which related genes are arranged in tandem, likely an evolutionary result of gene duplication events followed by strong natural selection (Dutheil et al., 2016; Kamper et al., 2006; Schirawski et al., 2010). Transcriptomic studies have shown that the expression of these genes is co-regulated. Clustered genes are upregulated upon host infection whereas flanking genes are not (Kamper et al., 2006; Lanver et al., 2018). In addition, clustered effectors are known to contribute to virulence, although with some degree of redundancy. Individual deletion of seven of these clusters lead to reduced or apathogenic phenotypes (Kamper et al., 2006; Schirawski et al., 2010). However, besides establishing a general role in virulence, reverse genetic studies have not been successful in establishing the function of effector gene clusters.

The plant immune system recognizes two major classes of pathogen-derived molecules, pathogen/microbial associated molecular patterns (PAMPs/MAMPs) and effectors. PAMPs are highly conserved, essential structural components of the pathogen and cannot be modified without large fitness costs. These include bacterial flagellin, fungal chitin, and β-glucans (Dodds & Rathjen, 2010; Jones & Dangl, 2006). The recognition of PAMPs is mediated by a large set of plasma membrane receptors called pattern recognition receptors (PRRs). PAMP binding to PRRs leads to a series of defense reactions that include the rapid production of ROS by membrane-anchored NADPH oxidases (RBOH proteins), increase in cytosolic Ca^2+^ levels, MAP kinase activation, and transcriptional reprogramming. Together, these responses are called PAMP-triggered Immunity (PTI). Consequentially, these signaling cascades are often manipulated by bacterial effectors in order to promote virulence (Couto & Zipfel, 2016). Likewise, the *U. maydis* effector Pep1 inhibits the activity of host apoplastic peroxidases involved in the production of ROS (Doehlemann et al., 2009; Hemetsberger, Herrberger, Zechmann, Hillmer, & Doehlemann, 2012).

The ubiquitin proteasome system is a regulatory mechanism that also controls multiple layers of the plant immune system, including PAMP recognition and downstream signaling. Particularly, E3 ubiquitin ligases (E3s) have been shown to regulate the activity and turnover of components involved in immune signaling. In *Arabidopsis thaliana* (hereafter Arabidopsis), the three closely related E3s, PUB22, PUB23, and PUB24 negatively regulate immunity by dampening PAMP recognition (Trujillo, Ichimura, Casais, & Shirasu, 2008). In contrast, the “*Arabidopsis TOXICOS EN LEVADURA*” gene family encodes RING domain E3s which are induced by PAMP treatment and whose mutation makes plants more susceptible to pathogens (Martínez-García, Garcidueñas-Piña, & P., 1996; Ramonell et al., 2005). E3s are also frequently targeted by effectors. For example, the *P. infestans* Avr3a effector stabilizes CMPG, an E3 whose degradation is necessary for immune reactions (Bos et al., 2010). Most strikingly, the *Pseudomonas syringae* effector AvrPtoB is an E3. It mediates the ubiquitination of several PRRs, leading to their degradation and thereby promoting virulence (Rosebrock, Zeng, Brady, Abramovitch, & Xiao, 2007).

Here we present the functional characterization of a major *U. maydis* effector cluster, the *pleiades* / cluster 10A, containing 10 putatively secreted proteins. By expressing these proteins in the plant cell, we show that 8 of the pleiades effectors inhibit PAMP-triggered immunity, thereby identifying a PTI-suppressive function of the cluster. We demonstrate that neofunctionalization followed gene duplication in the case of the two paralogous effectors Taygeta1 (Tay1) and Merope1 (Mer1). Finally, we provide evidence that Mer1 targets and modifies the activity of RFI2 homologs, a conserved family of RING E3s that are involved in early immune responses and control of flowering time in plants. Arabidopsis plants expressing Mer1 show decreased immunity as well as early flowering.

## Results

### The *pleiades*, a cluster of secreted proteins that contribute to virulence

The *U. maydis pleiades* cluster (Cluster 10A) encodes ten proteins, which, beyond their predicted secretion signals, lack any sequence similarity to known protein domains. Additionally, no cysteine residues (which are frequent in apoplastic effectors) are encoded outside of the predicted secretion signals. The *pleiades* contain three gene families, A (UMAG_03745, UMAG_03746, UMAG_03747 and UMAG_03750) B (UMAG_03748, UMAG_03749) and C (UMAG_03752, UMAG_037453), based on protein sequence similarity (25% or more). Two genes (UMAG_03744 and UMAG_03751) encode for proteins without homology to any other protein in the cluster (Fig1 a). While none of the *pleiades* show paralogs outside of the cluster, orthologs of all three gene families are well conserved across the sequenced smut fungi and display high synteny between *U. maydis*, *Sporisorium reilianum*, and *S. scitamineum*, Thus, the gene cluster is conserved among these species (Table S1).

**Figure 1.**
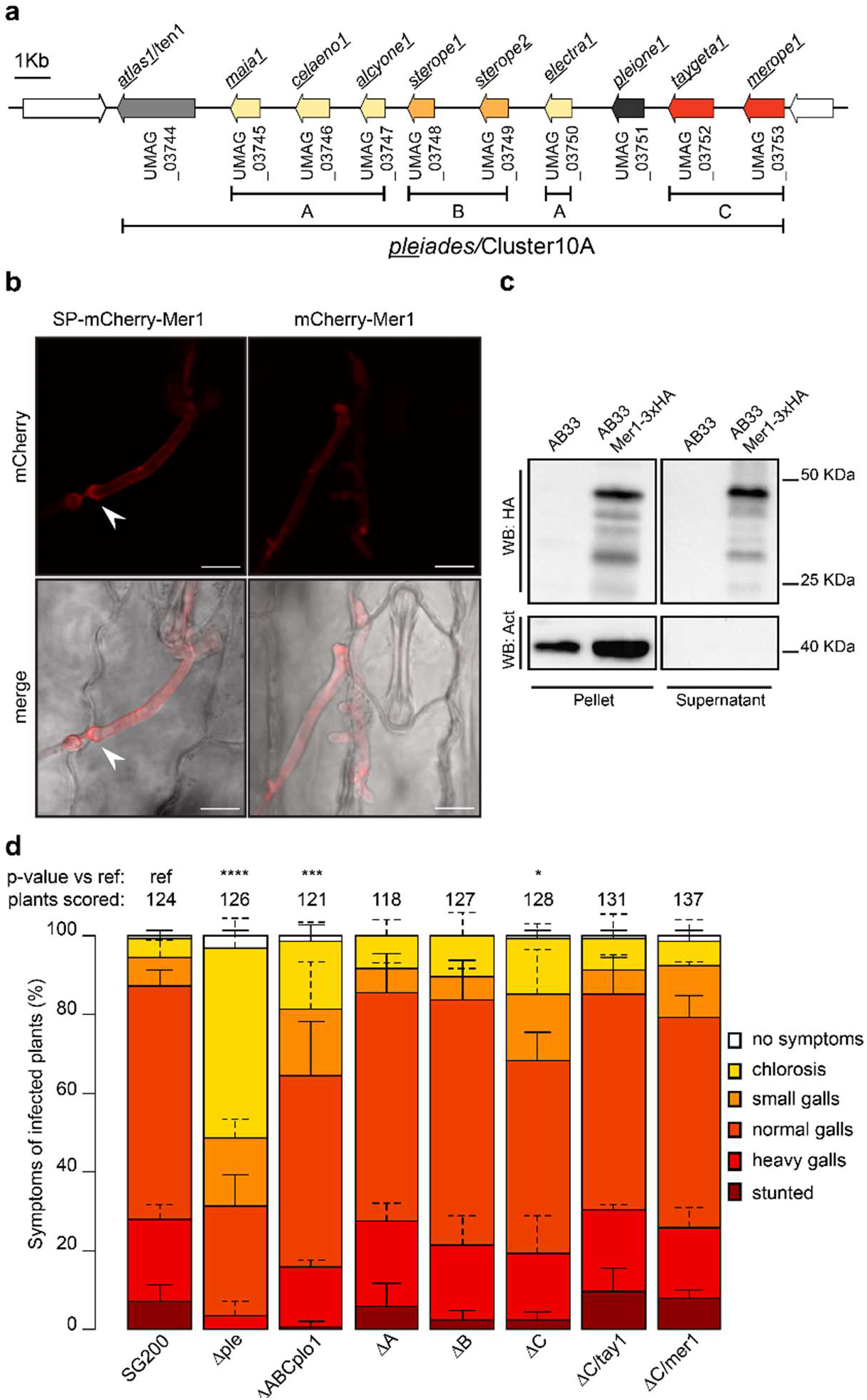
Pleiades, a cluster of secreted proteins contributes to the virulence of *U. maydis*. **a.** Schematic representation of the cluster 10A/Pleiades on chromosome 10 of *U. maydis*. Paralogous genes are represented with the same color and assigned to different families (A, B, C, respectively). Genes neighboring the cluster, which do not contain a predicted secretion signal, are shown in white. **b.** Secretion of Mer1 during maize infection. Left: plants infected with a *U. maydis* strain expressing SP_Mer1_-mCherry-Mer1 show mCherry signal mainly at the periphery of the hyphae and in the cell to cell crossings (arrowhead). Right: plants infected with a strain expressing mCherry-Mer1 (without its secretion signal) show a diffuse mCherry signal, without accumulation in the periphery of the hyphae. Upper panels: mCherry fluorescence, lower panels: bright field-mCherry merge. Scale bar = 10 μm. Pictures were taken 3 dpi. **c.** Secretion of Mer1 in axenic culture. SP-Mer1-3×HA was expressed in the *U. maydis* strain AB33 (which is able to filament *in vitro*). Total proteins were extracted from the pellet and secreted proteins were precipitated from the culture supernatant. The extracts were subjected to western blot with α HA or α Actin antibodies. Mer1-3×HA is detectable in the pellet and supernatant fractions whereas the cytosolic actin control is only detected in the pellet fraction. **d.** Disease symptom scoring of maize seedlings infected with *U. maydis*. SG200 (progenitor strain) or its derivative strains harboring complete or partial deletion of the *pleiades* cluster and their ectopic complementations were infected in seven-day old seedlings, and disease symptoms were rated 12 dpi. Data represent mean ± SD from three independent experiments, n= total number of scored plants. Significant differences between strains were analyzed by the Fisher’s exact test with Benjamini-Hochberg correction for multiple comparisons (*p<0.05, ** p<0.01, *** p<0.001, **** p<0.0001).

The transcriptional regulation of the *pleiades* during the various life-stages of *U. maydis* reveals that they are transcriptionally upregulated during the biotrophic phase of the fungus (Lanver et al., 2018). Analysis of the Pleiades with SignalP 5.0 (Almagro Armenteros et al., 2019) predicted the presence of secretion signals for all proteins with high confidence, whereas analysis of the two immediately neighboring genes (UMAG_03743 and UMAG_03754), showed that they code for proteins without secretion signals (Table S2). We verified the secretion of two Pleiades by monitoring the localization of mCherry fusions of Tay1 (UMAG_03752) and Mer1 (UMAG_03753) proteins expressed by *U. maydis* during biotrophic growth in maize. Three to four days post infection (dpi), *U. maydis* expressing Mer1_SP_-mCherry-Mer1 or Tay1_SP_-mCherry-Tay1 showed localization of the mCherry signal in the edges and tips of the hyphae (Fig 1b, Fig supplement S1c) and at the host cell to cell crossings (Fig 1b, arrowheads). On the other hand, *U. maydis* expressing mCherry-Mer1 (without its predicted secretion signal) showed a diffuse localization of the mCherry signal throughout the whole hyphae (Fig 1b). We next used the *U. maydis* AB33 strain to express Mer1 and Tay1 in axenic culture. AB33 filaments *in vitro* in response to nitrate, mimicking to some degree developmental changes induced during host colonization (Brachmann, Weinzierl, Kämper, & Kahmann, 2001). Using the strong, constitutive *otef* promoter we found that full length Mer1-3×HA and Tay1-3×HA accumulated in both, the cell pellet and culture supernatant fractions, whereas the non-secreted protein Actin was only detectable in the cell pellet fraction (Fig 1c, Fig supplement S1d). When using the *tay1* promoter (which is ten times stronger than the *mer1* promoter (Lanver et al., 2018), we could not detect the expression of these proteins in vitro (Fig 1d). Taken together, the presented data shows that Mer1 and Tay1 are soluble proteins secreted by *U. maydis* into the biotrophic interphase upon host colonization.

Deletion of the whole *pleiades* cluster has been shown to impair virulence of *U. maydis* (Kamper et al., 2006). Therefore, we generated deletions strains in the solopathogenic SG200 *U. maydis* background to dissect the specific virulence contribution of individual gene families within the *pleiades*. We generated deletions of the entire gene cluster, individual gene families or all the genes in the cluster except *atl1* (formerly *ten1*, UMAG_03744), since the latter was previously reported to contribute to virulence (Erchinger, 2017). Deletion of the whole cluster had the strongest effect on virulence. Plants infected with this strain predominantly showed mild disease symptoms like chlorosis and small galls (Fig 1d). Simultaneous deletion of the gene families A, B, C together with *plo1* also showed a considerable reduction in virulence, although not as strong as in the whole cluster deletion. Finally, deletion of family C showed a mild defect in virulence, which could be complemented ectopically by either *tay1* or *mer1*. Deletion of family A or B alone did not show any measurable effect on virulence (Fig 1d). Altogether our experiments show that the *pleiades* contribute to virulence additively and that *atl1* has the greatest impact.

### Pleiades suppress early defense responses

To investigate how the *pleiades* contribute to virulence, we analyzed early host defense responses upon infection with *U. maydis* strain SG200 or its derivative SG200 *Δple*. One of the first signaling and defense responses that plants activate upon recognition of invading microbes is the accumulation of reactive oxygen species (ROS) in the apoplastic space, a process that is usually suppressed by effectors from virulent pathogens (Dodds & Rathjen, 2010; Jones & Dangl, 2006). We assessed the production of ROS at the infection sites by staining plants 36 hours post infection with diamino-benzidine (DAB), which forms a brown precipitate in the presence of H_2_O_2_ (Y.-H. Liu, Offler, & Ruan, 2014; Thordal-Christensen, Zhang, Wei, & Collinge David, 1997) and examined them by wide field and fluorescence microscopy. Maize leaves infected with SG200 *Δple* showed strong DAB precipitation that accumulated around the invading hyphae, whereas SG200-infected leaves were hardly stained, and hyphae appeared mostly transparent by wide field microscopy (Fig 2a). Hyphae that did not show DAB precipitation could be stained by the chitin-binding WGA-Alexa Fluor 488 dye (WGA-AF488), indicating that all examined areas were colonized by *U. maydis* (Fig 2a). These results show that *pleiades* contribute to the suppression of ROS upon maize colonization by *U. maydis*.

**Figure 2.**
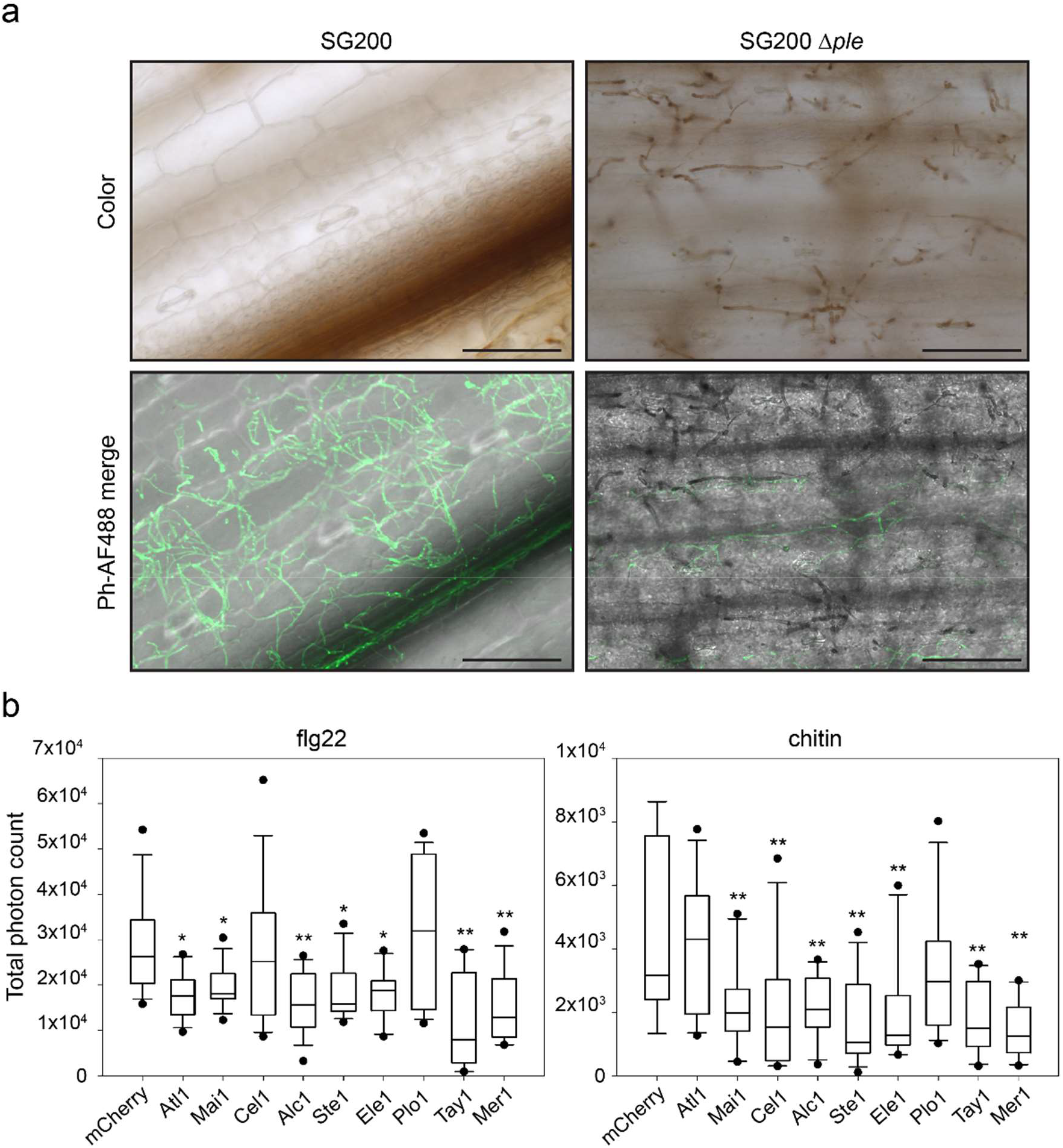
Pleiades contribute to the suppression of early defense responses. **a.** Accumulation of H_2_0_2_ in infected leaves visualized by diamino benzidine (DAB) staining. Left panels: *U. maydis* SG200 colonizes the host tissue without showing strong DAB precipitation. Right panels: *U. maydis* SG200 *Δple* shows pronounced precipitation of DAB around the hyphae upon colonization of the host. Upper panels: color pictures showing the brown DAB precipitate. Lower panels: Phase contrast/AF488 merge. WGA-AF488 (which binds chitin) was used as a counter stain to show the presence of hyphae not stained by DAB, since the precipitate covers and mask chitin from being bound by WGA. Samples were collected 36h post infection. Scale bar = 100 μm. Notice that the color pictures and the Phase contrast/Fluorescent pictures are taken with different cameras, therefore the fields of the upper and lower panels don’t overlay perfectly. **b.** PAMP-triggered ROS burst in *N. benthamiana.* Transient expression of *pleiades* leads to reduction of the oxidative burst triggered by flg22 (left) or chitin (right). Plants expressing mCherry were used as the reference control. Total photon counts over 40 (flg22) or 30 minutes (chitin) are shown as box plots. Data is a pool of three (flg22) or four (chitin) independent experiments, n= 15 or 12 plants respectively. Significant differences between proteins were analyzed by ANOVA with Benjamini-Hochberg correction for multiple comparisons (* p<0.05, ** p<0.01).

To complement H_2_O_2_ visualization in the *U. maydis*-maize pathosystem and to test whether the Pleiades have a direct impact on the early PTI response, we expressed each of the proteins (without their predicted secretion signal) in *Nicotiana benthamiana* and tested their ability to suppress PAMP-triggered ROS production. Leaf disks of plants expressing each of the *pleiades* were treated with the PAMPs flg22 or chitin and ROS production was monitored over 30-40 minutes using a luminol-based assay (Lloyd, Schoonbeek, Martin Trick, Zipfel, & Ridout, 2014). The ROS-burst response in plants expressing mCherry cloned in the same vector used for expression of the *pleiades* was used as reference control. Strikingly, all Pleiades except Plo1, were able to inhibit the PAMP-triggered ROS burst compared to the control (Fig 2b, Fig supplement S2). Most proteins inhibited the oxidative burst irrespective of the PAMP used, except Atl1 which was specific for flg22 and Cel1 which was specific for chitin. All ROS burst curves can be seen in the Supplementary figure S2. Taken together, our results indicate that Pleiades are secreted and likely translocated into the host cells where they inhibit the PAMP-triggered ROS burst. In addition, the *N. benthamiana* assays also imply that the molecular targets of Pleiades must be conserved across monocots and dicots.

Tay1 and Mer1 from family C had the strongest effect on PAMP-triggered ROS production in the used plant expression system, leading us to focus further experiments on these genes (Fig 2b, Fig supplement S2c).

### The paralogs Tay1 and Mer1 target different cellular compartments

To clarify how Tay1 and Mer1 suppress ROS production, we analyzed their sub-cellular localization. We constructed mCherry fusions of Tay1 and Mer1 and expressed them in maize epidermal cells by biolistic bombardment. GFP- nuclear localization signal (NLS) was co-transformed and used as a nuclear marker. Confocal microscopy showed that mCherry-Tay1_28-398_ localized primarily to the cytoplasm and almost no mCherry signal was detected in the nucleus (Fig 3a). On the other hand, mCherry-Mer1_23-341_ was in the nucleus as well as the in the cytoplasm (Fig 3a). These results lead us to hypothesize that these paralogs may target different cellular compartments in order to suppress ROS production. We tested this hypothesis using a mis-localization approach. We fused Tay1 and Mer1 to either an NLS, NES (nuclear export signal) or Myr (myristoyl lipid anchor signal, targeting proteins to the cytoplasmic side of the plasma membrane) and expressed these proteins in *N. benthamiana*. Myc-tag fusions of Tay1_28-398_ and Mer1_23-341_ were used as controls since this tag is not expected to affect cellular localization. Plants were treated with flg22 and ROS production was monitored over time as described above. Localizing each paralog to different cellular compartments had opposite effects. Tay1 showed an increased ROS-burst inhibition upon forced cytoplasmic localization (NES and Myr fussions). Fusing Tay1_28-398_ to the NLS signal did not affect its activity (Fig 3b). In contrast, Mer1 showed decreased ROS-burst inhibition upon forced cytoplasmic localization (NES and Myr fussions). Fusing Mer1_23-341_ to the NLS signal did not affect its inhibitory activity (Fig 3c). Hence, by integrating the maize microscopy studies with the *N. benthamiana* mis-localization data, we propose that Tay1 acts in the host cytoplasm whereas Mer1 acts in the host nucleus.

**Figure 3.**
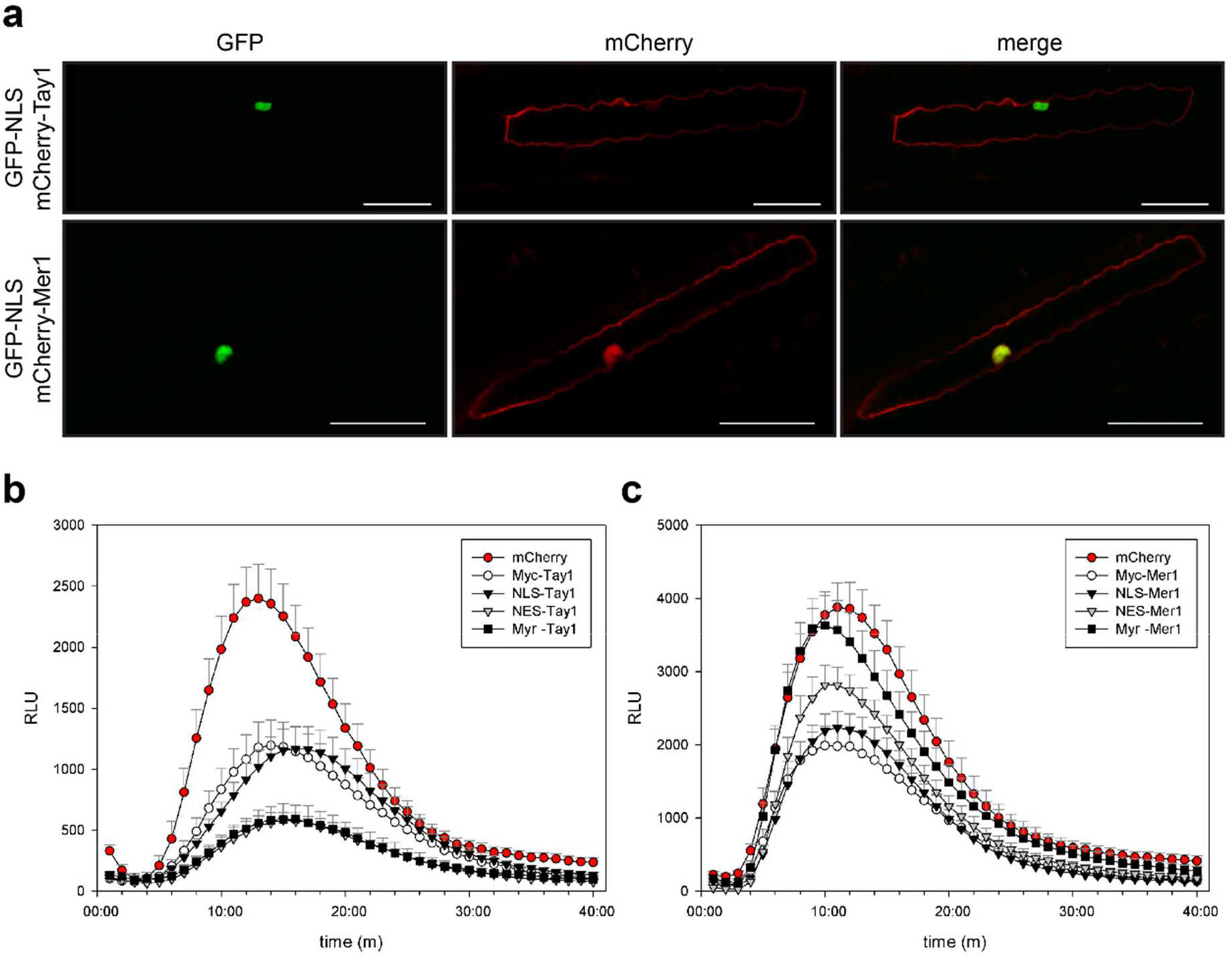
Tay1 and Mer1 act in different cell compartments. **a.** mCherry-Tay1 and mCherry-Mer1 were co-expressed with GFP-NLS in maize epidermal cells by biolistic bombardment. mCherry-Tay1 (upper panels) localizes preferentially to the cytoplasm whereas mCherry-Mer1 (lower panels) localizes to the cytoplasm and nucleus. Scale bar = 50 μm. **b. c.** Sub-cellular localization affects the ROS burst inhibiting activity of Tay1 and Mer1 differentially. Plants expressing mCherry were used as reference control showing maximum ROS-burst and plants expressing Myc-Tay1 (b) or Myc-Mer1 (c) were used to monitor their intrinsic ROS-burst inhibitory activity without miss localization. Curves show flg22-triggered ROS burst over 40 minutes. **b.** Removing Tay1 from the nucleus (NES or Myr) increases its inhibitory activity, whereas targeting Mer1 to the nucleus (NLS) does not modify its inhibitory activity. **c** Removing Mer1 from the nucleus (NES or Myr) decreases its inhibitory activity, whereas targeting Mer1 to the nucleus (NLS) does not modify its inhibitory activity. Data, mean ± SEM, is a pool of three independent experiments, n= 15. Only positive error bars are shown for clarity.

### Mer1 interacts with RFI2, a family of E3-Ligases

To identify host targets of Tay1 and Mer1 we performed yeast two-hybrid (Y2H) screens against a cDNA library from *U. maydis*-infected maize tissues. Whereas the screen with a binding domain fusion with Tay1 (BD-Tay1_28-398_) did not lead to the identification of any reproducible interactors, screening with BD-Mer1_23-341_ led to the identification of 72 clones growing on high stringency media (-L, -W, -H, A). Of these, 31 corresponded to proteins with homology to the Arabidopsis RING domain, E3 red and far red insensitive 2, *RFI2* (Chen & Ni, 2006b). To independently verify this interaction, we cloned four of the five predicted RFI2 homologs from maize, the two predicted homologs from Arabidopsis (we renamed the only characterized homolog At*RFI2A* and its uncharacterized paralog *RFI2*B) and the only predicted homolog from *N. benthamiana* into a prey vector and tested their interaction against BD-Mer1 _23-341_. All homologs showed interaction with BD-Mer1_23-341_ on intermediate stringency media (−L, -W, -H). Additionally, the homologs ZmRFI2A, ZmRFI2B, ZmRFI2T, and NbRFI2 showed interaction on high stringency media (Fig 4a). ZmRFI2T is a truncated variant, isolated from the cDNA library, that lacks the RING domain (Fig supplement S4a). Its interaction with Mer1 implies that Mer1 does not require the RING domain for interaction with the RFI2 homologs.

**Figure 4.**
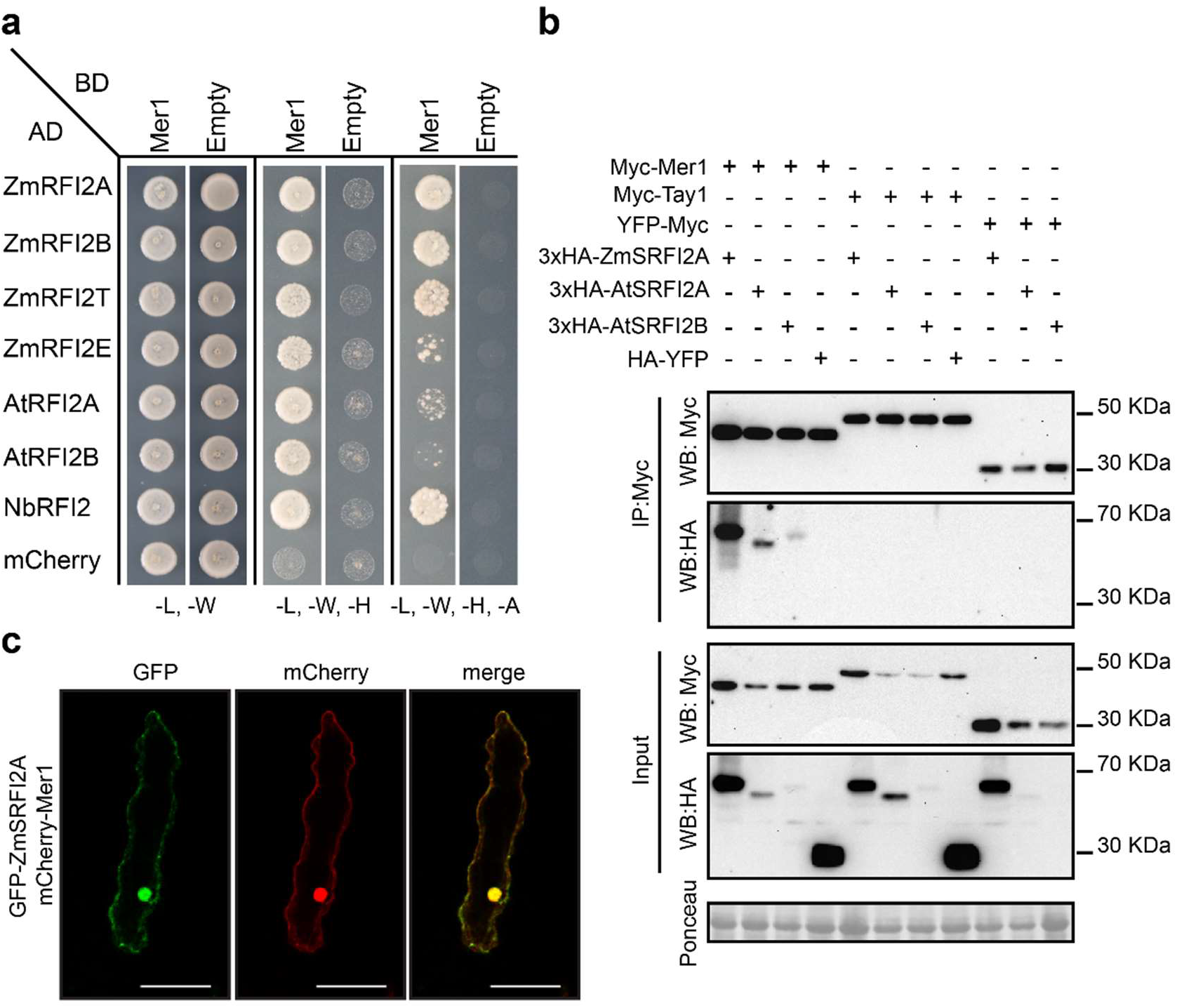
Mer1 interacts with members of the RFI2 family in the plant nucleus. **a.** Y2H assay showing interaction of Mer1 with RFI2 homologs from maize, Arabidopsis and *N. benthamiana*. Yeast strains were grown on SD medium lacking indicated amino acids/nucleotides. Growth in media lacking leu (L) and trp (W) is used as transformation control. Growth in media lacking leu (L), trp (W) and his (H) or leu (L), trp (W), his (H) and ade (A) indicates protein interaction. **b.** RFI2 proteins co-immunoprecipitate with Mer1 *in planta*. Proteins were co-expressed in *N. benthamiana*, total proteins were immunoprecipitated with α Myc magnetic beads (IP: Myc) and blotted with specific antibodies. 3×HA-ZmRFI2Am, 3×HA-AtRFI2Am and 3×HA-AtRFI2Bm co-immunoprecipitate with Myc-Mer1 specifically. Protein loading is indicated by Ponceau staining in the input fraction. **c.** mCherry-Mer1 and GFP-ZmRFI2Am co-localize in the plant nucleus. Proteins were transiently co-expressed in the epidermis of *N. benthamiana* by *Agrobacterium* mediated transformation. Scale bar = 50 μm.

To further confirm the interaction between Mer1 and RFI2 homologs we performed co-immunoprecipitation assays (Co-IPs). Since RING E3s are known to auto-ubiquitinate, thus mediating their own degradation *in vivo* (Carvalho, Saraiva, Maia, Abreu, & Duque, 2012; Stone et al., 2005), we constructed stabilized versions containing alanine substitutions in the Zn-coordinating residues of the RING domain (Fig supplement S4b) which we named “stabilized RFI2” (SRFI2). 3×HA-ZmSRFI2A, 3×HA-AtSRFI2A, or 3×HA-SAtRFI2B were co-expressed with Myc- Mer1_23-341_ in *N. benthamiana*. Total proteins were extracted and incubated with α-Myc magnetic beads. All three RFI2 homologs tested were able to co-immunoprecipitate in the presence of Myc-Mer1_23-341_ but not in the presence of Myc-Tay1_28-398_ or YFP-Myc (negative controls, Fig 4b). Additionally, HA-YFP was not co-precipitated in the presence of Myc-Mer1_23-341_ (Fig 4b). These results show that the interaction between Mer1 and RFI2 homologs is specific.

Since we established earlier that Mer1 targets the host nucleus (Fig 3), we tested whether RFI2 homologs also localize to this cellular compartment. As before, we co-bombarded fluorescently labeled proteins into the maize epidermis and verified their localization by confocal microscopy. GFP-ZmSRFI2A co-localized with mCherry-Mer1_23-341_ in the nucleus and cytoplasm. GFP-ZmSRFI2A and mCherry-Mer1_23-341_ co-localized in the nucleus and to a lesser extent, in the cytosol. (Fig 4c). Co-expression of GFP-ZmSRFI2A, GFP-AtSRFI2A or GFP-AtSRFI2B with mCherry-Mer1_23-341_ in the epidermis of *N. benthamiana* showed similar results, although the nuclear localization of the E3s was more pronounced in this system (Fig supplement S4c). Taken together, our results show that Mer1 specifically interacts with members of the RFI2 E3 family and that this interaction likely happens in the nucleus. The interaction with maize, *N. benthamiana*, and Arabidopsis homologs also confirms our earlier assumption that the Pleiades’ targets are conserved across monocots and dicots.

### Mer1 modifies the ubiquitination activity of RFI2 homologs

In Arabidopsis*, RFI2*A has been linked to seedling de-etiolation responses and photoperiodic flowering. *rfi2*A plants show early flowering under long day conditions (Chen & Ni, 2006a, 2006b). Additionally, the rice homolog *APIP6*, has been linked to immunity (C.-H. Park et al., 2012). To clarify the role of *rfi2* homologs in immunity we took advantage of the genetic tractability of Arabidopsis. We generated plants expressing Mer1_23-341_ and compared them to the *rfi2A*, *rfi2B*, and *rfi2A/rfi2B* double knock outs. As expected, Arabidopsis plants expressing Myc-Mer1_23-341_ showed a reduced PAMP-triggered ROS burst compared to wild-type (Col-0) plants (Fig 5a). Interestingly, this phenotype occurred even at low expression levels of Mer1, indicating that the mechanism of ROS-burst suppression is requiring only minor effector amounts (Fig supplement S5c). The *rfi2A* and *rfi2B* knock out plants also showed a reduced ROS burst, similar to that of Myc-Mer1_23-341_ plants. Finally, the double knocks out *rfi2A*/*rfi2B* did not show a further ROS burst reduction upon flg22 treatment compared to the single knock outs (Fig 5b). This data, together with evidence from the literature, indicate that RFI2 family proteins are the targets of Mer1. They are necessary for ROS production in response to PAMPs and this role is conserved across monocots and dicots.

**Figure 5.**
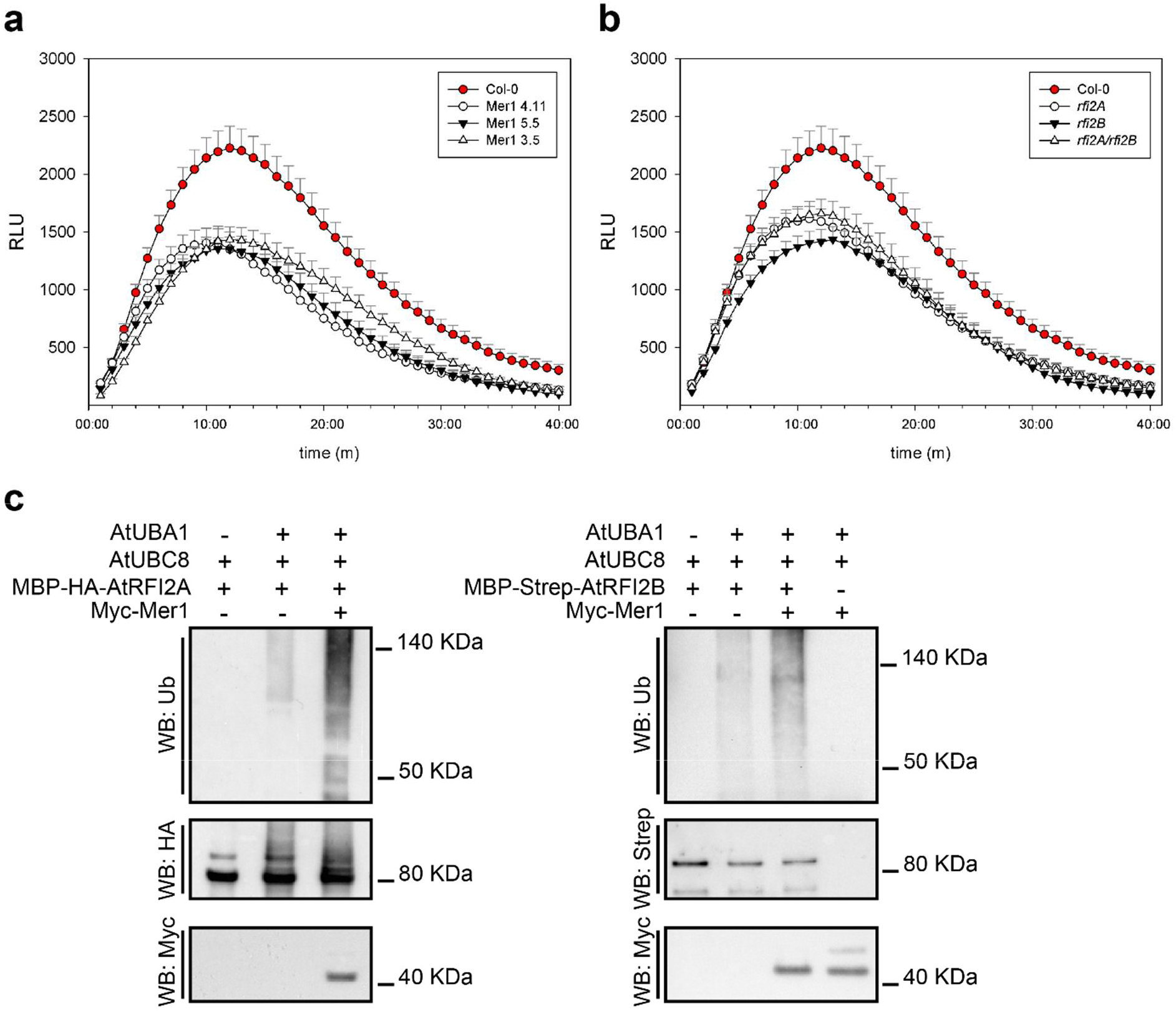
Mer1 affects the activity of AtRFI2s. **a.** flg22-triggered ROS burst in *Arabidopsis*. Myc-Mer1 lines show reduced ROS burst upon flg22-treatment compared to their Col-0 background. **b.** Both *rfi2* homologs of *Arabidopsis* contribute to the flg22-triggered ROS burst response. Either single mutant or the double mutant show reduced oxidative burst compared to their Col-0 background. Only positive error bars are shown for clarity. Notice that all the measurements were done simultaneously, panels a and b are shown separately only for clarity. Data, mean ± SEM, is a pool of four independent experiment, n= 20. **c.** Mer1 promotes the autoubiquitination activity AtRFI2s *in vitro*. Proteins were affinity-purified from *E. coli*, used to perform ubiquitination reactions and the results were assayed by blotting with specific antibodies. Addition of Mer1 promotes the ubiquitination of AtRFI2A (left) or AtRFI2B (right). Reactions excluding E1 or E3s were used as negative controls.

Arabidopsis plants expressing Mer1_23-341_ also showed an early flowering phenotype compared to their Col-0 background when grown under long day conditions (Fig supplement S5a, S5b). In contrast, *rfi2A* and *rfi2B* plants did not flower, while the *rfi2A/rfi2B* double knockout flowered slightly late (Fig supplement S5b).

Since expression of Mer1_23-341_ phenocopied the *rfi2* knockouts, we hypothesized that Mer1 must have an inhibitory effect on the E3s. To test this hypothesis, we produced and purified Myc-Mer1_23-341_, MBP-HA-AtRFI2A and MBP-Strep-AtRFI2B from *Escherichia coli* and assayed the effect of Mer1 on the ubiquitination activity of the E3s *in vitro*. Both E3s showed a moderate auto-ubiquitination activity, that was strongly enhanced by addition of Mer1 (Fig 5c). In the case of AtRFI2A, we could detect the higher molecular weight autoubiquitination products of the E3 by western blot directly with α-HA and α-Ubiquitin antibodies. In the case of AtRFI2B, we could only detect the ubiquitination products with the α-Ubiquitin antibody. As negative controls, we performed reactions lacking either E1 or E3 enzymes, which showed no ubiquitination products.

Taken together, our results indicate that the effector Mer1 targets the RFI2 family of E3 ubiquitin ligases which mediate PAMP-triggered ROS burst. Mer1 inhibits their activity by promoting auto-ubiquitination, likely leading to higher degradation of the E3s and decreased PAMP-triggered ROS-burst.

## Discussion

The analysis of the *U. maydis* genome revealed that a significant number of putative effector genes, whose expression is induced upon host infection, are physically clustered in the genome (Kamper et al., 2006). Similar to prokaryotic operons, gene clusters are commonly found in diverse fungi to co-regulate functionally connected genes (Howlett, Idnurm, & Heitman, 2007; Keller & Hohn, 1997; Rokas, Wisecaver, & Lind, 2018). As no functional characterization of an *U. maydis* effector cluster has been reported, a common role for these clusters during biotrophy was based on assumptions (Kamper et al., 2006). Here, by using an approach analogous to those used for fungal metabolic gene clusters (Howlett et al., 2007), although relying on heterologous gene expression rather than knock outs, we demonstrate that eight clustered effectors share the ability to suppress PAMP-triggered immunity independent of their sequence relationship. Therefore, for the *pleiades*, their position in the genome is a better predictor of their function than sequence homology. This finding raises the question of whether other clustered effectors also share redundant functions. Moreover, we increased our understanding of effector functions in plant pathogenic fungi. Particularly in smuts, only six effectors have been functionally characterized (Djamei et al., 2011; Hemetsberger et al., 2012; Lay-Sun Ma, 2018; Mueller, Ziemann, Treitschke, Assmann, & Doehlemann, 2013; Redkar et al., 2015; Tanaka et al., 2014), and only three are likely translocated and function within the host cytosol (Djamei et al., 2011; Redkar et al., 2015; Tanaka et al., 2014). Our results and hypothesis on how the *pleiades* affect the interaction between *U. maydis* and maize are summarized in Fig 6.

**Figure 6.**
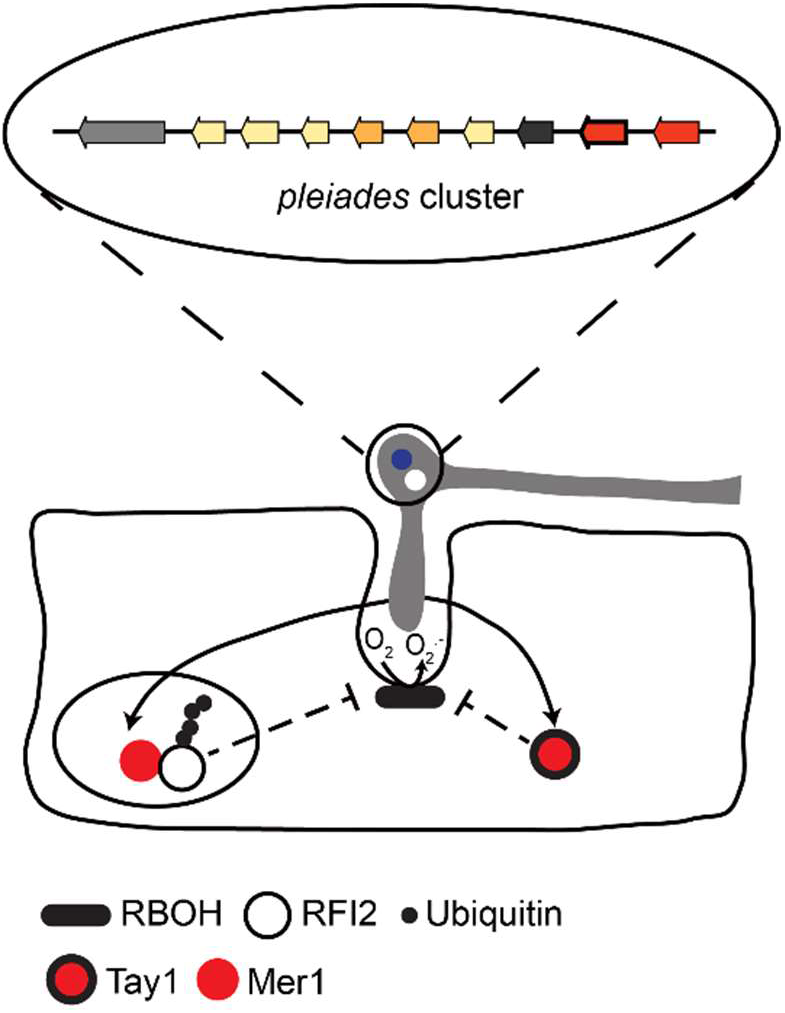
Infection of maize by *U. maydis* triggers RBOH-mediated production of reactive oxygen species (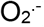) around invading hyphae. To counteract this early defense response, *U. maydis* secretes the Pleiades, a heterogeneous group of translocated effectors whose genes are clustered in the genome. Most Pleiades show inhibition of PAMP-induced ROS-burst. Mer1, targets the host nucleus, where it binds to and promotes auto-ubiquitination of RFI2s, a family of E3s that control the production 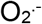. The Mer1 paralog, Tay1, inhibits the production of 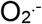 from the cytoplasm.

Although we have not demonstrated translocation of the Pleiades into host cells directly, our data strongly supports that the Pleiades are symplastic effectors. First, none of the mature Pleiades have cysteine residues, considered a hallmark of apoplastic effectors. Disulfide bond formation occurs under the oxidative conditions of the apoplast, stabilizing protein structures (Lanver et al., 2017; Win et al., 2012). Additionally, the fact that infected plants show higher H_2_O_2_ levels around hyphae of the *Δple* mutant than around the progenitor strain agrees with the suppression of PAMP-triggered ROS-burst upon expression of the Pleiades in the plant cytosol. Moreover, we have found that Mer1 targets and modifies the auto-ubiquitination of RFI2 homologs, a conserved family of E3s that function within the plant nucleus to control ROS production. This finding is consistent with the concept that effectors tend to target hubs of the immune system (Mukhtar et al., 2011; Weßling et al., 2014). The effector AvrPiz-t from *Magnaporthe oryzae* was shown to be translocated into rice cells, where it targets a member of the RFI2 family, APIP6 (C.-H. Park et al., 2012). The fact that two effectors from unrelated pathogens target the same E3 highlights its importance as an essential component for host defense. In an independent study, it has been shown shown by immunogold-labelling and electron microscopy that one of the Pleiades, Atl1/Ten1, is translocated into maize cells during infection by *U. maydis* (Erchinger, 2017). The same study further suggests that Atl1 interacts with the PP2C phosphatase ZmPP26. PP2C phosphatases have been shown to be regulators of immunity (Couto & Zipfel, 2016; C.-J. Park et al., 2008; Schweighofer et al., 2007). Thus, our findings together with previous studies, suggest that all Pleiades are likely translocated effectors that act in the host cytosol. The mechanisms or signals that lead to translocation of fungal effectors into host cells are still under debate, and for *U. maydis* remain completely unknown (Ribot et al., 2013; Tanaka et al., 2015; Wawra et al., 2013). The identification of a critical mass of experimentally validated translocated effectors will help to elucidate commonalities of translocated effectors that could be used for predicting their translocation into host cells based on sequence features.

Using heterologous plant systems to study effector function introduces the problem that host-specific factors might be missed. On the other hand, effectors that target highly conserved proteins and pathways can be identified. Bearing this in mind, most members of the Pleiades must target highly conserved components of the PAMP-triggered immunity in plants, which we have demonstrated for Mer1. Plo1 on the other hand, which did not suppress ROS-burst in *N. benthamiana*, might target a maize specific protein or pathway. Alternatively, it might have not been functional upon expression in in *N. benthamiana* or may simply have a different function. Nonetheless, we propose that the Pleiades are valuable tools to identify and dissect conserved elements of the plant immune system.

Our work provides a prime example of functional redundancy by demonstrating that eight clustered effectors collectively inhibit PAMP-triggered ROS-burst. Where the pathogen relies on a battery of tools to ensure that critical defense responses are efficiently suppressed during host colonization. This redundancy might allow subpopulations of *U. maydis* to lose some of the *pleiades* in order to avoid recognition by particular host genotypes, while still maintaining the ability to suppress PAMP-triggered ROS burst (Sánchez-Vallet et al., 2018). Alternatively, there might be some degree of specialization amongst the Pleiades. For example, different Pleiades may vary in their efficiency at targeting different host protein alleles. Maintaining many different Pleiades could therefore be an adaptation to a genetically diverse host such as maize (Schnable et al., 2009). Indeed, some of the Pleiades show a certain degree of functional and mechanistic specialization. Cel1 and Atl1 inhibit ROS burst only when triggered by chitin or flg22, respectively. In the case of Atl1, the fact that a fungal effector specifically inhibits flg22 dependent immunity might be an indication that it targets BAK1-dependent pathways. This co-receptor interacts with many PRRs and thus is necessary for the perception of many PAMPs, including flg22 but not chitin (Cao et al., 2014; Heese et al., 2007; Smakowska-Luzan et al., 2018). Further support for functional specialization of the *pleiades* comes from the analysis of their expression. (Skibbe, Doehlemann, Fernandes, & V., 2010) found that most pleiades are upregulated to a higher degree during seedling infection compared to flowers or mature leaves, whereas *plo1, tay1*, and *mer1* are equally upregulated during seedling and flower infection. Additionally, during seedling infection, the expression of *atl1*, *mai1, ste1*, and *ele1* peak 2 days after infection while the rest of the *pleiades* peak 4 days after infection or later (Lanver et al., 2018).

Considering that in smut fungi effectors frequently emerge by gene duplication followed by rapid diversification (Dutheil et al., 2016; Schirawski et al., 2010), it is likely that a similar evolutionary history favored the appearance of new functional variants within the *pleiades* cluster. For the paralogs Tay1 and Mer1, which are encoded by adjacent genes in the cluster and share 31% identity at the amino acid level, our mis-localization studies show that they suppress ROS production in different sub-cellular compartments (cytoplasm or nucleus). Furthermore, *tay1* is widespread across smuts whereas *mer1* is restricted to the closely related *U. maydis*, *S. relianum*, and *S. scitamineum*, where *tay1* and *mer1* are neighboring genes (Fig supplement S1a). Thus, *tay1* and *mer1* show the characteristics of a neofunctionalization event.

Finally, the phenotypes of Arabidopsis plants expressing Mer1, reduced immunity and early flowering, suggest a connection between both processes. Although a similar link has been reported before for the Arabidopsis *pub13* and rice *spl11* knock outs, these phenotypes have been attributed to increased salicylic acid (SA) levels in these mutants. However, since high SA levels are known to increase PAMP triggered immunity (J. Liu et al., 2012; Tateda et al., 2014), it is unlikely that the early flowering phenotype of Mer1 plants is due to increased levels of this hormone. In addition, although at*rfi2A* has been shown to negatively regulate flowering time in the Arabidopsis accession Ws-0 (Chen & Ni, 2006b) neither At*rfi2A*, At*rfi2B*, nor the At*rfi2A/*At*rfi2B* double mutant showed an early flowering phenotype in the Col-0 accession. However, these two accessions are notoriously different with regards to their flowering behavior (Passardi et al., 2007), which might indicate the presence of an additional factor that needs to be inactivated in Col-0 in order to reveal the early flowering phenotype. Accordingly, this factor could be affected by Mer1. Nonetheless, an effector that dampens immunity while simultaneously promoting flowering would be a great advantage for smuts, which usually sporulate only in the host floral tissues. This is in line with the concept that effectors frequently target regulatory nodes to shift the balance from immunity to growth and development (Uhse & Djamei, 2018).

In conclusion, we have shown that Pleiades, a heterogeneous group of proteins, whose genes cluster in the genome of *U. maydis*, share the ability to suppress PAMP triggered immunity. a conserved link between PAMP triggered defense and flowering time in plants, therefore making the interception point of both pathways an excellent target for manipulation by pathogens. Considering the strong evidence provided, that Pleiades act through different mechanisms makes the other Pleiades valuable starting points to identify conserved and possibly novel players of PAMP-triggered immunity in plants

## Materials and methods

### Gene accession numbers

*atlas*1: UMAG_03744, *maia*1: UMAG_03745, *celaeno*1: UMAG_03746, *alcyone*1: UMAG_03747, *sterope*1: UMAG_03748, *sterope*2: UMAG_03749, *electra*1: UMAG_03750, *pleione*1: UMAG_03751, *taygeta*1: UMAG_03752, *merope*1: UMAG_03753, Zm*rfi2*A: Zm00001d037596, Zm*rfi2*B: Zm00001d009837, Zm*rfi2T*: Zm00001d040789, Zm*rfi2*E: Zm00001d052697, At*rfi2*A: at2g47700, At*rfi2*B: at3g05545, Nb*rfi2*: Niben101Scf00369g11013.1.

### Protein sequence analysis

For candidate effector genes, sequences were screened for the presence of a secretion signal with SignalP 5.0 (Almagro Armenteros et al., 2019). Protein alignments were done using CLC Main Workbench 7.7.2.

### Plasmids, cloning procedures and generation of *U. maydis* strains

All plasmids were generated by standard molecular procedures (Sambrook, Russell, & Sambrook, 2006). *E. coli* Mach1 (Thermo Fisher Scientific, Waltham, MS, USA) was used for all DNA manipulations. Plasmids used for expression of “untagged” effector proteins in *N. benthamiana* (Figure 2b) were generated by Gateway Cloning (Katzen, 2007). All other plasmids were generated by the GreenGate system (Lampropoulos et al., 2013). The modules used were either amplified by PCR or obtained from the published system. Additionally, we generated two GreenGate destination vectors. pECGG, is based on a pET backbone and was used for expression of proteins in *E. coli*. pADGG, is based on a pGAD backbone and was used as prey vector for yeast two hybrid assays.

*U. maydis* knock out strains were generated by homologous recombination with PCR-derived constructs (Kämper, 2004). For complementations and protein expression, strains were generated by insertion of p123 derivatives into the *ip* locus (Loubradou, Brachmann, Feldbrugge, & Kahmann, 2001). Transformants were verified by southern blot and/or PCR. All plasmids and strains used in this study can be found in Supplementary materials.

### Maize infection assays

Pathogenicity assays and disease symptom scoring were performed as described by (Kamper et al., 2006). Briefly, *U. maydis* SG200 and its derivatives were cultured in liquid YepsLight (0.4% yeast extract,0.4% peptone and 2% sucrose) at 28°C to an OD_600nm_ of 0.6-0.8. Cells were pelleted by centrifugation at 2400 g for 10 m and resuspended in H_2_O to an O.D._600nm_ of 1. The suspensions were then syringe-inoculated in seven-day old maize seedlings (variety Early Golden Bantam, Old Seeds, Madison, WI, USA). Maize was grown in a temperature-controlled glasshouse (14h light/10h dark, 28°C/20°C) and disease symptoms were assessed 12 dpi. Filamentous growth of the *U. maydis* strains was tested by spotting the respective strains in potato dextrose agar containing 1% activated charcoal. The experiments were repeated at least 3 times.

### Transient expression in maize by biolistic bombardment

Biolistic bombardment was performed according to (Djamei et al., 2011) with minor modifications. Briefly, 1.6 μm gold particles were coated with plasmid DNA encoding the indicated constructs under the CaMV35S promoter. Bombardment was performed on seven-day old maize leaves cv B73 using a PDS-1000/HeTM instrument (BioRad) at 900 p.s.i. in a 27 Hg vacuum. Fluorescence was observed by confocal microscopy 18-24h after transformation. The experiments were repeated at least 3 times.

### Confocal microscopy

Confocal microscopy was performed with a Zeiss LSM 700 confocal microscope. GFP was excited at 488 nm using an argon laser. Fluorescence emission was collected between 500-540 nm. mCherry was exited at 561nm and emission was collected between 578-648 nm. Images were processed with ZEN blue 2.3 lite.

### Protein secretion in *U. maydis*

Detection of secreted proteins from fungal cultures was performed as described in (Djamei et al., 2011). Briefly, *U. maydis* was grown in CM medium until the culture reached an O.D._600nm_ of 0.6-0.8, centrifuged, resuspended in AM medium and incubated for 6h to induce filamentation. Cultures were centrifuged to separate supernatant and pellet fractions. Supernatant proteins were precipitated with 10% trichloroacetic acid and 0.02% sodium deoxycholate (final concentration) and resuspended in 100 mM Tris pH 8. Total proteins from the cell pellet were extracted by directly adding SDS loading buffer and, a spatula tip of glass beads and incubating samples in a vortex for 10 m. Protein extracts from the cell pellets and culture supernatants were subjected to immunoblotting using α-HA (Sigma-Aldrich, St. Louis, MO, USA) and α-Actin antibodies (Invitrogen, Waltham, MA, USA). Effector proteins were detected with α-HA antibody. Actin was used as a lysis control. *In vitro* secretion experiments were repeated 3 times.

For visualization of secreted proteins in infected tissues, maize plants were harvested 3-4 dpi infection and analyzed by confocal microscopy as described above. All strains used for protein secretion assays in maize carried constructs driven by the Um*tay1* promoter. Experiments were repeated at least 3 times.

### DAB-WGA staining in infected maize tissue

H_2_O_2_ production was detected in infected maize tissues using diamino-benzidine (DAB) (Sigma-Aldrich, St. Louis, MO, USA). Plants infected with *U. maydis* were harvested 36 h post infection. The 3^rd^ leaf was removed, cut with a scalpel and the bottom part was dipped in DAB solution (1mg/ml, pH 3.8) in the dark at room temperature for at least 16 h. Leaves where de-stained by several washes with ethanol/chloroform (4:1) until chlorophyll was no longer visible and stored at 4 **°**C until further processing. Samples were washed with PBS and fungal chitin was stained with a solution of wheatgerm agglutinin coupled to AlexaFluor488 (WGA-AF488, Invitrogen) (WGA-AF488 10μg/ml, Tween20 0.02% in PBS) by applying vacuum 3 times. Visualization of the stained samples was performed by direct observation with a widefield microscope equipped with Apotome2 (Axio Imager.Z2 sCMOS camera, Axiocam colour camera and Apotome2). DAB was visualized with a color camera or by Phase contrast. Chitin, labeled with wheatgerm agglutinin AlexaFluor488, was visualized by Apotome2 structured illumination with a 480/40nm excitation filter and 525/50nm emission filters. Images were processed with ZEN blue 2.3 lite. The experiments were repeated at least 3 times.

### Yeast transformation and two-hybrid assays

All yeast protocols were done according to the Yeast Protocols Handbook (Clontech, Mountainview, CA) with minor modifications. Strain AH109 was transformed with bait vectors (pGBKT7 and derivatives) and strain Y187 was transformed with prey vectors (pAD, pADGG and derivatives) by the LiAc/PEG method. All positive clones were verified for the presence of the corresponding plasmid by DNA extraction with 20mM NaOH and PCR (Supplementary materials).

The Y2H screen was performed by mating the strain carrying pGBKT7-Mer1_23-341_ (bait) against a yeast library carrying cDNA from maize tissue infected with *U. maydis* in the pAD vector (Farfsing, 2004). Positive clones were selected in high stringency media (SD -leu, -trp, -ade, -his).

For one to one matings AH109 pGBKT7-Mer1_23-341_ or AH109 pGBKT7 were mated against Y187 pADGG-ZmRFI2A, Y187 pADGG-ZmRFI2B, Y187 pADGG-ZmRFI2T, Y187 pADGG-ZmRFI2E, Y187 pADGG-AtRFI2A, Y187 pADGG-AtRFI2B, Y187 pADGG-NbRFI2A or Y187 pADGG-mCherry. Diploid cells were selected in SD –leu, -trp. For plate drop out assays, diploid cells were grown in liquid SD -leu, -trp o.n., Cells were pelleted by centrifugation at 500 g for 3 m and resuspended in sterile H_2_O to an O.D._600nm_ of 1. Serial dilutions were made in H_2_O and 5 μl of the suspensions were plated in SD -leu, -trp (growth control), SD -leu, -trp -his (intermediate stringency) and SD -leu, -trp, -his, -ade (high stringency). Growth in intermediate or high stringency media 4 days post inoculation indicated positive interactions. All plate drop out assays were repeated 2 times.

### Plant growth conditions, Arabidopsis lines and flowering time experiments

*N. benthamiana* and *A. thaliana* plants were grown in controlled short-day conditions (8h light/16h dark, 21°C) in Einheitserde as substrate. The plants were watered by flooding for 15 min every two days. At*rfi2*A is: SAIL_1222_B08, At*rfi2*B is: SALK_089110C, *rfi2*A/*rfi2*B double knock out was created by crossing and screening for the double knock out from the F2 populations (Supplementary materials). Mer1 Arabidopsis plants were created by floral dipping using the following construct: 35S:Myc-Mer1_23-341_ (Supplementary materials). For flowering time experiments *A. thaliana* plants were stratified for 3 days at 4°C in the dark and grown in the same substrate as above in long day conditions (16h light/8h dark, 21°C/16°C). Cool white fluorescent lamps were used as the light source (64-99 *μ*mol m−2 sec−1). Flowering time was determined as the total number of rosette leaves plus the cauline leaves when the stem was 1 cm long. Flowering time experiments were repeated at least 3 times.

### Protein production in *N. benthamiana* and Co-immunoprecipitation

For *in vivo* Co-IP assays, *A. tumefaciens* GV3101 (pSoup) carrying the expression constructs were grown o.n. in LB supplemented with the appropriate antibiotics at 28°C, centrifuged for 10 min at 1600g, resuspended in ARM buffer (*Agrobacterium* resuspension medium, 10mM MES-NaOH pH 5.6, 10mM MgCl_2_, 150 μM Acetosyringone) to an OD_600nm_ of 0.2 and incubated for 3h at RT. Cultures carrying the appropriate constructs were then mixed 1:1 and infiltrated in *N. benthamiana* with a needless syringe. Plants were incubated for 60h, frozen in liquid N_2_ and total proteins were extracted from 450mg of tissue in 2ml IP buffer: HEPES 50mM pH7.5, NaCl 100mM, Glycerol 10%, EDTA 1mM, Triton X-100 0.1%, PMSF 1mM and 1 protease inhibitor table/50ml (Roche cOmplete EDTA free cat# 05056489001). Extracts were cleared by centrifugation 10 m at 20000g speed 3 times. Proteins were immunoprecipitated by adding 30ul of α-c-Myc magnetic beads (μMACS Anti-c-myc Miltenyi Biotec, cat# 130-091-284) and incubated for 2h at 4°C with rotation. Samples were washed 4 times with 300 μl IP buffer and proteins were eluted by adding 50 μl of 2× SDS loading buffer at 95°C. 10-15 μl of the extracts were analyzed by SDS-PAGE followed by Western blot with α-c-Myc (Sigma-Aldrich, St. Louis, MO, USA) or α-HA (Sigma-Aldrich, St. Louis, MO, USA) antibodies. Experiments were repeated 2 times.

### ROS burst assays in *N. benthamiana* and *A. thaliana*

5-6-week-old *A. thaliana* or 4-5-week-old *N. benthamiana* plants were used for the assay. *N. benthamiana* was infiltrated with *A. tumefaciens* carrying the appropriate constructs resuspended in ARM buffer to a final OD_600nm_ of 0.1 and incubated for 48h. Leaf disks (4mm) were cut and floated in water o.n. Water was removed, and elicitors were added. flg22 elicitation solution consisted of Horseradish peroxidase (HRP 10 μg/ml, Sigma-Aldrich cat# P6782), L-012 (34ug/ml Fujifilm WAKO cat# 120-04891) and flg22 (100nM) in H_2_O. Chitin elicitation solution was prepared as follows: 50mg of chitin (Sigma-Aldrich cat# C9752) were ground with mortar and pestle in 5 ml of H_2_O for 5 m, transferred to a falcon tube, microwaved for 40 s, sonicated for 5 m, centrifuged at 1800 g for 5 m, the supernatant was transferred to a new tube, vortexed for 15 m and stored at 4**°**C. Before use, the suspension was diluted 1:1 in H_2_O and supplemented with HRP (as described above) and luminol (34 μg/ml Sigma-Aldrich cat# 123072). Reactive oxygen species production was monitored by luminescence over 30-40 minutes in a microplate reader (Synergy H1, BioTek). At least 3 plants per construct/genotype were used in each experiment. All experiments were performed at least 3 times.

### Protein production in *E. coli and* ubiquitination assays

Recombinant protein production was done as follows: His-AtUBA1 and His-AtUBC8 were produced in *E. coli* strain BL21 AI (ThermoFisher). All cultures were grown in LB Ampicillin at 37**°**C to an OD_600nm_ of 1, cold shocked on ice for 20m and protein production was induced by adding L-Arabinose (0.2%), or L-Arabinose plus ethanol (1.5%) in the case of UBA1. Cultures were further incubated for 5h at 22**°**C and cell pellets were frozen at −80**°**C until further processing. All remaining proteins (His-Myc-Mer1_23-341_, His-MBP-HA-AtRFI2A, His-MBP-Strep-AtRFI2B) were produced in *E. coli* BL21 pLys. Cultures were grown in LB Spectinomycin at 37**°**C to an OD_600nm_ of 0.6 and protein production was induced by adding IPTG (0.5 mM). Cultures were further incubated 3h at 37**°**C or o.n. at 22**°**C in the case of His-Myc-Mer1_23-341_ and cell pellets were frozen at −80**°**C until further processing. The version of Mer1 used here was codon optimized for *E. coli*.

All proteins were extracted in: Tris/HCl 50mM pH 7.5, NaCl 300mM, Glycerol 10%, imidazole 20 mM and protease inhibitors as described above. Proteins were purified with Ni-NTA agarose resin (Quiagen, cat# 30210), washed with extraction buffer supplemented with imidazole 160 mM and eluted with the same buffer supplemented with imidazole 250-400mM. Proteins Were concentrated and imidazole removed by centrifuging the extracts through a 30 KDa cutoff filter (Vivaspin 20 centrifugal concentrator, Sigma-Aldrich cat# Z614602-12EA). Proteins were stored in 50% glycerol at −80 **°**C until further processing.

Ubiquitination reactions were done in 45 μl as follows: Tris pH 7.5 50 mM, MgCl_2_ 5mM, KCl 25mM, ZnCl_2_ 50 μM, DTT 0.25 mM, ATP 5mM, Ubiquitin 3 μg (Sigma-Aldrich, cat# U6253), AtUBA1 100 ng, AtUBC8 250 ng, AtRFI2A 250 ng, AtRFI2B 250 ng, Mer1 250 ng. Reactions were incubated at 30°C for 2 h and stopped by adding 15 ul of 4X SDS loading buffer and incubated at 95**°**C for 5 m. Analysis of the reactions was done by SDS-PAGE followed by by western blotting with StrepTactin-HRP Conjugate (Bio Rad, cat # 1610380), α-c-Myc (Sigma-Aldrich, St. Louis, MO, USA), α-HA (Sigma-Aldrich, St. Louis, MO, USA) and α-Ubiquitin (P4D1, Abcam cat# ab139101) antibodies. Experiments were repeated at least 3 times.

### Statistical analyses

Maize infection assays were analyzed by the Fisher exact test in R, as described by (Stirnberg & Djamei, 2016). All other statistical analyses were performed with GraphPad Prism 8.0. ROS burst data was analyzed by ANOVA with Benjamini-Hochberg correction for false discovery rate. Flowering time data was analyzed by ANOVA, Tukeys. Statistical significance was evaluated at the level of p<0.05.

---

**Table S1:**
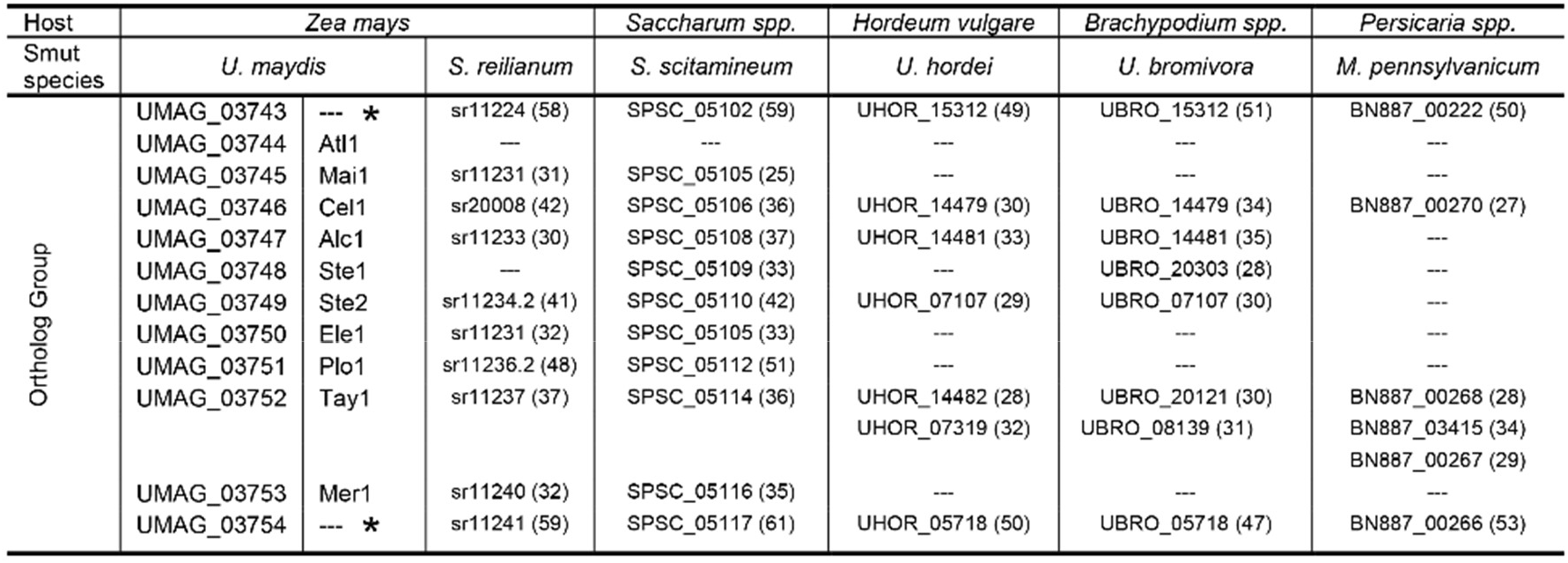
Conversation of Pleides protiens across different smuts

**Table S2:**
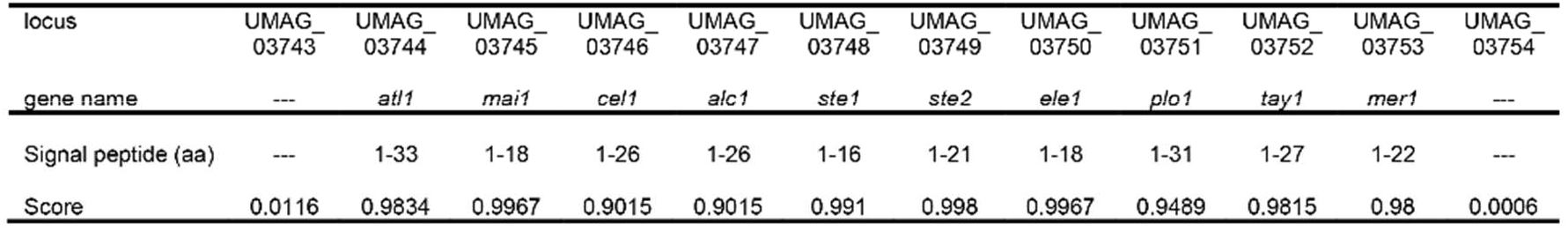
Signal peptide predictions

**Figure 1_ Figure supplement 1.**
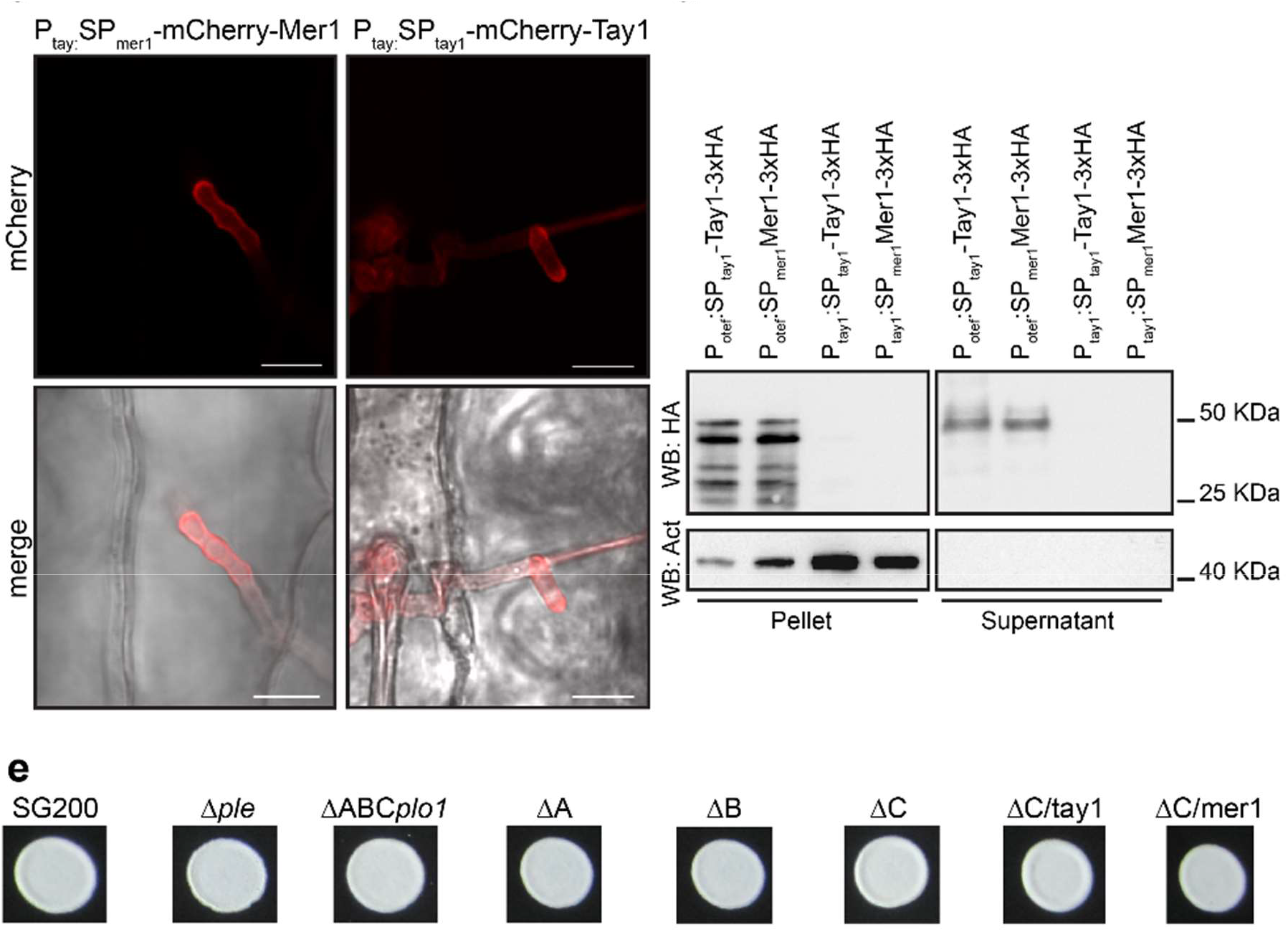
**a.** Conservation of the Pleiades proteins across different smut species. For each *U. maydis* protein, a list of orthologs is shown, the percentage of similarity is shown in brackets. Absence of an ortholog in any given genome is indicated by “---”. Genes neighboring the Pleiades cluster (which do not contain a predicted secretion signal) are marked by “*”. **b.** Prediction of signal peptides across the *U. maydis* Pleiades. Signal peptides were predicted with SignalP-5.0. The table shows the secretion peptides in amino acids and the likelihood of the prediction (1 being the highest). Lack of a secretion signal is depicted by “---”. **c.** Secretion of Tay1 and Mer1 during maize infection. Left: Plant infected with the *U. maydis* strain SG200 ∆*C* carrying the construct P_tay1_:SP_Tay1_-mCherry-Tay1 in the *ip* locus. Right: Plant infected with the *U. maydis* strain SG200 ∆*C* carrying the construct P_tay1_:SP_Mer1_-mCherry-Mer1 in the *ip* locus. Notice the mCherry signal accumulation at the periphery and tip of the hyphae. Upper panels: mCherry fluorescence, lower panels: bright field-mCherry merge. Scale bar = 10 μm. Pictures were taken 3 dpi. **d.** Secretion of Tay1 and Mer1 in axenic culture. Constructs harboring SP_tay1_-Tay1-3×HA or SP_Mer1_-Mer1-3×HA were expressed in the strain AB33 under the *tay1* or *otef* promoter. Total proteins were extracted from the pellet and secreted proteins were precipitated from the culture supernatant. The extracts were subjected to western blot with α HA or α Actin antibodies. Tay1-3×HA and Mer1-3×HA could be detected in the pellet and supernatant fractions only when the expression was driven by the strong synthetic *otef* promoter. Actin could only be detected in the pellet fraction. **e.** Filamentation of *U. maydis* strains used in Fig1d. Strains were spotted on PD-charcoal plates and pictures were taken 24h after, showing that all strains retain the ability to filament, a prerequisite for infection.

**Figure 2_ Figure supplement 1.**
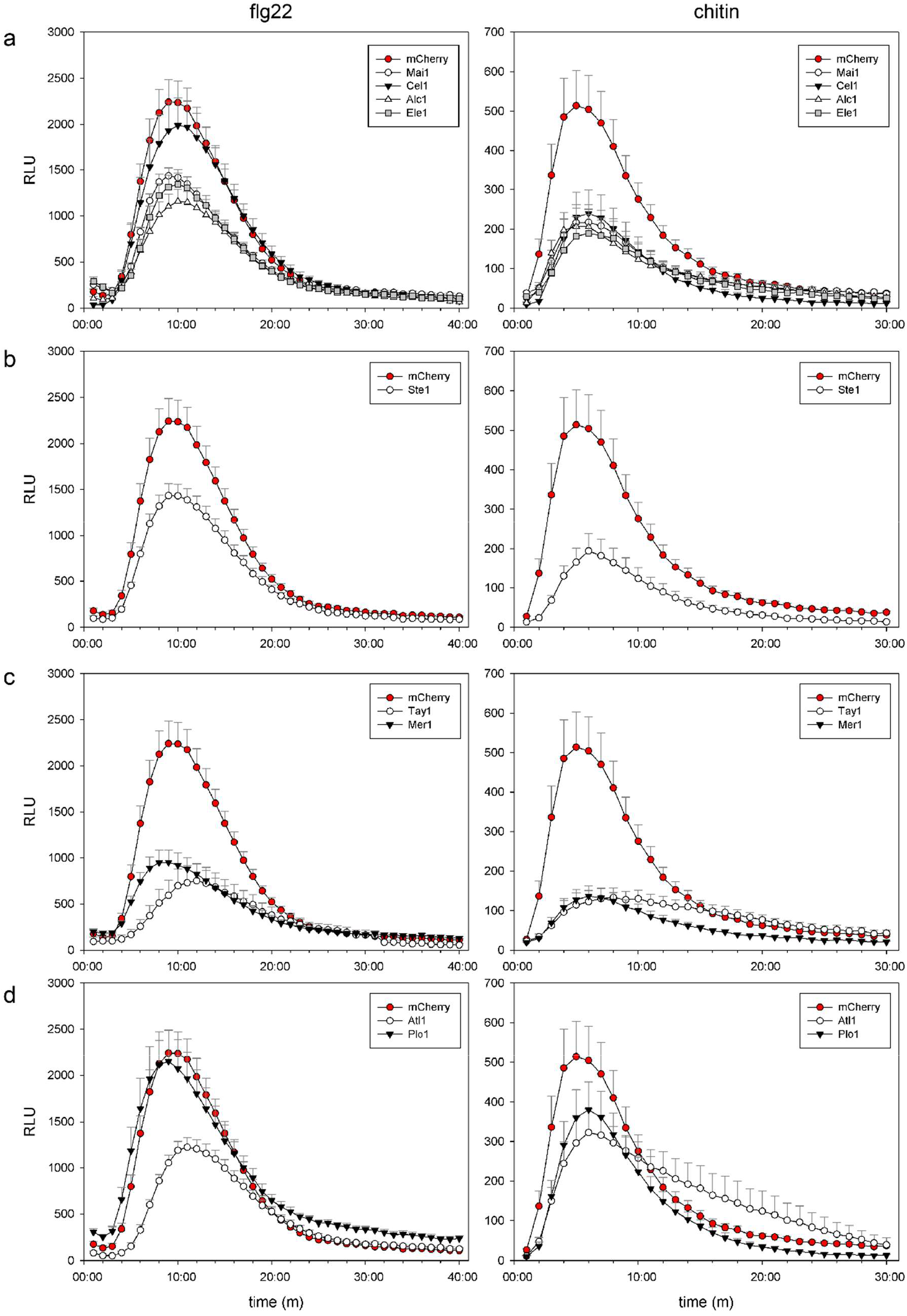
PAMP-triggered ROS burst curves corresponding to Fig2b. Left: flg22, right: chitin. **a.** Curves corresponding to proteins from family A. **b.** Curves corresponding to proteins from family B. **c.** Curves corresponding to proteins from family C. **d.** Curves corresponding to Atl1 and Ple1. Data is the mean ± SEM. Only the positive error bar is shown for clarity. Notice that all the measurements were done simultaneously, curves were split according to protein family only for clarity.

**Figure 4_ Figure supplement 1.**
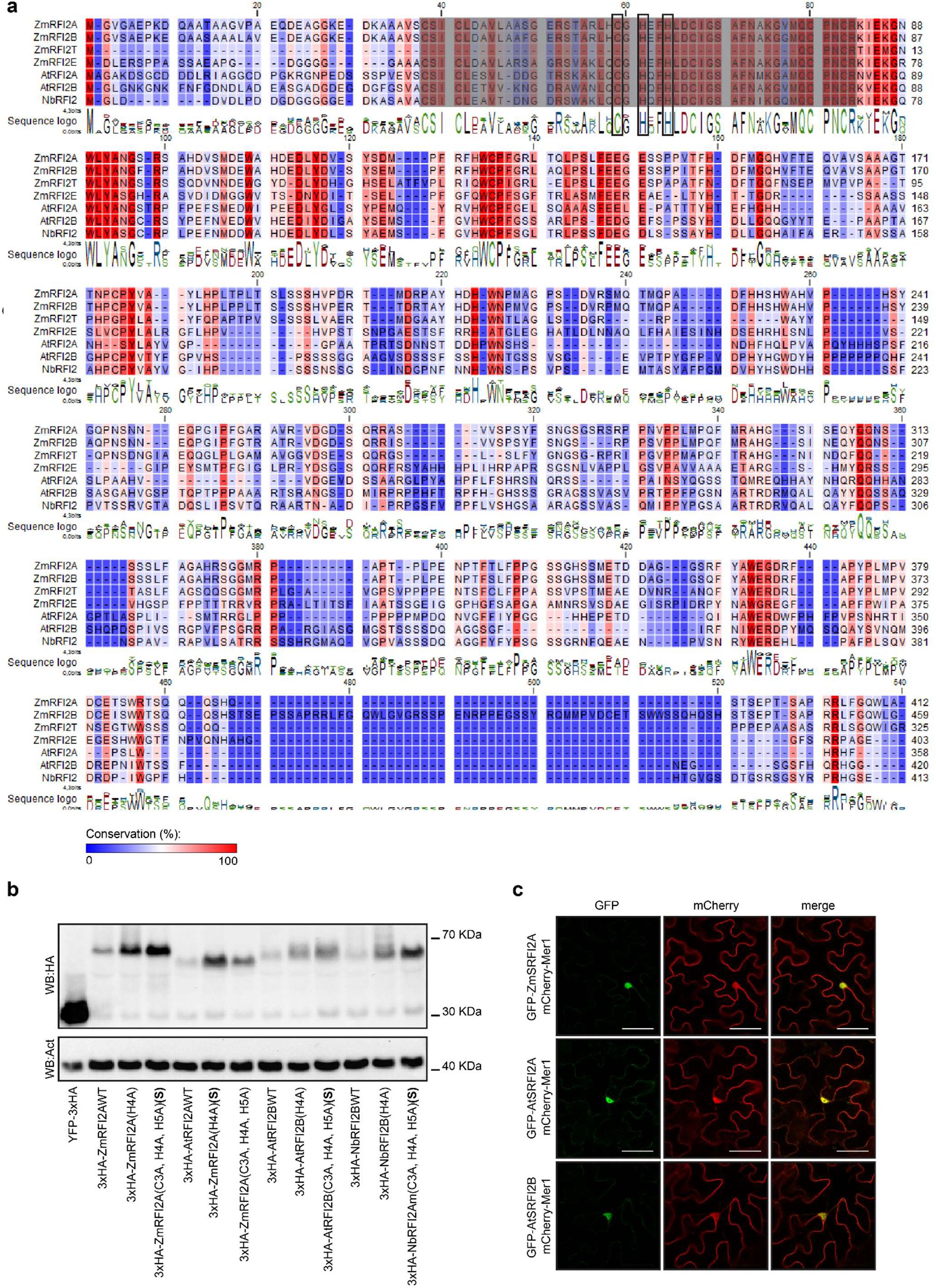
RFI2 is a family of E3 ubiquitin ligases whose localization is enriched in the nucleus. **a.** Protein alignment of selected members of the RFI2 family. Background color indicates conservation (red: 100%, blue 0%). Sequences correspond to the proteins used in Fig 4a. The residues shaded within the alignment mark the RING domain. Within the RING domain, Zn-coordinating residues 3, 4 and 5 are shaded. **b.** Mutation of the Zn-coordinating residues stabilizes RFI2s. Western blot α HA showing the expression of WT, single A substitutions, or triple A substitutions of the shaded residues in part a. For each homolog, the most stable mutant was named “S”. YFP-3×HA is shown for comparative reason. Western blot α Actin was used as loading control. **c.** Co-localization of Mer1 with different members of the RFI2 family in the epidermis of *N. benthamiana*. Scale bar = 50 μm.

**Figure 5_ Figure supplement 1.**
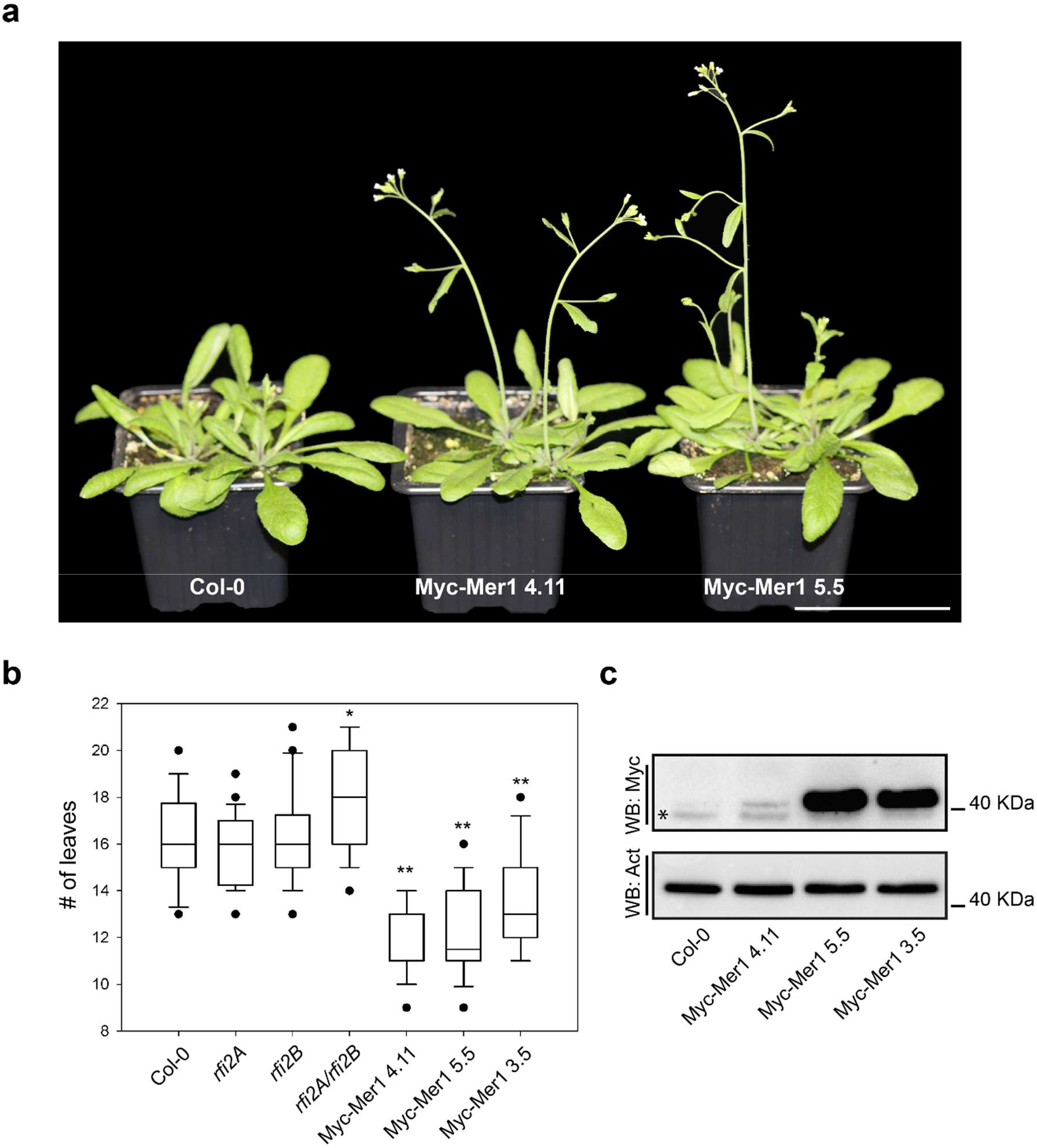
Mer1 *Arabidopsis* plants flower earlier than *wt* plants. **a.** Flowering of Col-0 and two independent Mer1 lines 29 days after planting. Plants were grown in long day conditions (16h light/8h dark). Scale bar = 6 cm. **b.** Quantification of flowering time, assessed as number of leaves at flowering day, is shown as box plots. Mer1 expressing plants show early flowering whereas the *rfi2A, rfi2B* and *rfi2A/rfi2B* Knockouts in the Col-0 background do not. One representative experiment is shown, n= 30 plants. Significant differences between lines were analyzed by ANOVA, Tukeys (* p<0.05, ** p<0.01). **c.** Expression of Mer1 in *A. thaliana*. Expression was assessed by western blot with α Myc antibodies, α Actin was used as loading control. “*” unspecific band.

## Acknowledgements

The research leading to these results has received funding from the European Research Council under the European Union’s Seventh Framework Programme ERC-2013-STG, Grant Agreement: 335691, the Austrian Science Fund (FWF): [P27429-B22, P27818-B22, I 3033-B22], and the Austrian Academy of Science (OEAW).

We would like to thank the GMI/IMBA/IMP core facilities for excellent technical support, especially, BioOptics, Molecular Biology Services and Plant Sciences Facility at Vienna BioCenter Core Facilities GmbH (VBCF). We are grateful to Mathias Madalinski for peptide synthesis, André Alcântara for useful comments on the manuscript and to Dr. J. Matthew Watson for proofreading and valuable feedback on the manuscript.

## Author Contribution

Conceptualization: Fernando Navarrete, Armin Djamei.

Funding acquisition: Armin Djamei.

Investigation: Fernando Navarrete, Nenad Grujic, Alexandra Stirnberg, David Aleksza, Hazem Adi.

Methodology: Fernando Navarrete, Marco Trujillo, Armin Djamei.

Project administration: Fernando Navarrete.

Resources: Fernando Navarrete, Alexandra Stirnberg, Nenad Grujic, Michelle Gallei, Janos Bindics, Marco Trujillo. Armin Djamei

Supervision: Armin Djamei.

Writing: Fernando Navarrete, Armin Djamei.

## Authors declaration

The authors declare that there is no conflict of interest in the research.

